# Resource supply drives even spacing of individuals along multiple trait axes in light-limited phytoplankton populations

**DOI:** 10.1101/338129

**Authors:** Simone Fontana, Mridul K. Thomas, Marta Reyes, Francesco Pomati

**Affiliations:** Department of Aquatic Ecology, Eawag, Swiss Federal Institute of Aquatic Science and Technology, Überlandstrasse 133, 8600 Dübendorf, Switzerland; Institute of Integrative Biology, Swiss Federal Institute of Technology (ETH), Universitätstrasse 16, 8092 Zürich, Switzerland

**Keywords:** phenotypic plasticity, trait evenness, niche complementarity, competitive ability, individual-level interactions

## Abstract

Individual-level variation arising from responses to environmental gradients influences population and community dynamics. How such responses empirically relate to the mechanisms that govern species coexistence is however not well understood. Previous results from lake phytoplankton communities suggested that the evenness of organisms in multidimensional trait space increases with resource limitation, possibly due to resource partitioning at the individual level. Here we experimentally tested the emergence of this pattern by growing two phytoplankton species *(Pseudokirchneriella subcapitata, Microcystis aeruginosa)* under a gradient of light intensity, in monoculture and jointly. Under low light (resource) conditions, the populations diversified into a wide range of phenotypes, which were evenly distributed in multidimensional trait space (defined by four pigment-related trait dimensions), confirming the observed field pattern. Our results provide prime experimental evidence that resource limitation induces even spacing of conspecific and heterospecific microbial phenotypes along trait axes, and advances our understanding of trait-based coexistence.

---

Many processes that are fundamental to community assembly, such as responses to abiotic gradients, direct species interactions (e.g. predation, parasitism), and competition (inter- as well as intraspecific), act at the level of individual organisms at the smallest temporal and spatial scales (Reiss *et al.* 2009; Clark *et al.* 2011). Stabilising mechanisms of coexistence explicitly require the relative importance of intra- vs. interspecific competition to be considered (Chesson 2000), and recent studies advocated the significant influence of individual variation in traits on community assembly and species coexistence (Clark 2010; Jung *et al.* 2010; Paine *et al.* 2011; Violle *et al.* 2012; De Laender *et al.* 2014; Barabás & D’Andrea 2016; Turcotte & Levine 2016).

Individual-level variation in traits is a common feature of all organisms, including clonal and microbial populations (Ackermann 2015). Resource limitation and fluctuations are an important driver of intraspecific phenotypic heterogeneity, also in microbial communities, where variation in phenotypes favours population fitness (Ackermann 2015; Schreiber *et al.* 2016; Zimmermann *et al.* 2018). For example, NH_4_+ limitation can trigger an increase in phenotypic heterogeneity in the N_2_-fixing bacterium *Klebsiella oxytoca,* with a benefit under resource fluctuations (Schreiber *et al.* 2016). A decrease in availability of resources like phosphorus and light in lakes was also shown to induce greater evenness (i.e. regularity) of individual phytoplankton cells in multidimensional trait space, possibly to minimise competition for limiting resources (Fontana *et al.* 2018). The latter community-wide pattern observed in nature invokes plasticity and/or selection inducing phenotypic changes that stabilise coexistence by allowing for resource partitioning among individuals and species. For instance, the relative composition and amount of pigments allows phytoplankton species to partition over the light spectrum, which is composed by different wavelengths that can be absorbed in different proportions depending on pigment-related traits (Stomp *et al.* 2004; Stomp *et al.* 2008).

In this study, we experimentally tested the hypothesis that limitation in resource supply, specifically light availability, induces phenotypic heterogeneity in terms of even spacing of individual phytoplankton cells across multiple photosynthesis-related traits, which can be measured by scanning flow-cytometry (SFC). Under competition for light, individual organisms should adjust pigment composition to capture different wavelengths of the light spectrum and reduce niche overlap, as has been observed at the species level (Stomp *et al.* 2004), and this pattern should emerge from both intra- and interspecific competition. Here we exposed two phytoplankton species belonging to different functional groups (the green alga *Pseudokirchneriella subcapitata* and the cyanobacterium *Microcystis aeruginosa*) to six different light intensities (Fig. 1), and quantified cell density and evenness of individual organisms in multidimensional trait space over time. The two species utilise the light spectrum in different ways: both possess chlorophyll (which captures light in the blue and red portions of the visible spectrum), while *M. aeruginosa* also possesses the accessory pigment phycocyanin (which captures light in the green-yellow portion, for the most part inaccessible to *P. subcapitata)* (Reynolds 2006). Monocultures were tested under direct light and under light passing through a culture of the competing species (hereafter indirect light). In this way, we simulated the effects of competition for light intensity and spectrum, while excluding direct interspecific interactions (such as through allelopathy). Additionally, we also tested a mixture of the two species (under direct light only) to account for direct individual-level interactions between species.

**Fig. 1.**
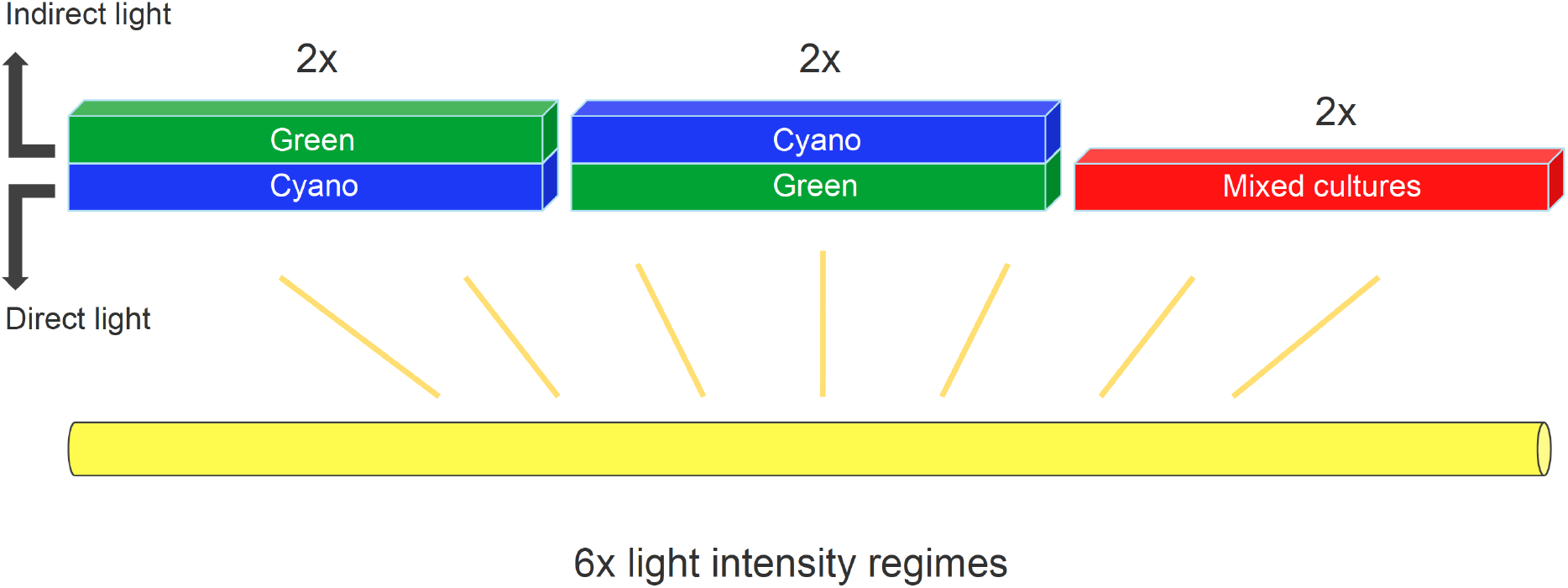
Illustration of the experimental design. For simplicity, only one out of two replicates and one out of six light treatments are reported. Green=*P. subcapitata*; Blue=*M. aeruginosa*; Red=Mixed cultures=*P. subcapitata+M. aeruginosa.*

If competitive interactions between individual organisms are the driving force of even spacing in acclimating phenotypes, we expect that the rate of change in trait evenness will range from positive to increasingly negative values as light intensity increases (Fig. 2). Under high levels of light, organisms will not minimise trait similarity by increasing evenness, but rather distribute freely in multidimensional trait space or tend towards few optimal trait combinations, leading to a decrease in trait evenness over time. Trait evenness has a maximum possible value (Fig. 2, dotted black line tending to a horizontal asymptote) and should reach a minimum as cell densities eventually induce resource limitation and competition, and therefore an increase in trait evenness (Fig. 2, blue and orange bands indicating a possible reversal of the curves). However, we did not expect the experiment to last long enough to observe a maximum or minimum in TED values. Our experiment zoomed in the initial trajectory of response, which can be approximated by a straight line (Fig. 2). Trait evenness values at carrying capacity should also be inversely proportional to light intensity.

**Fig. 2.**
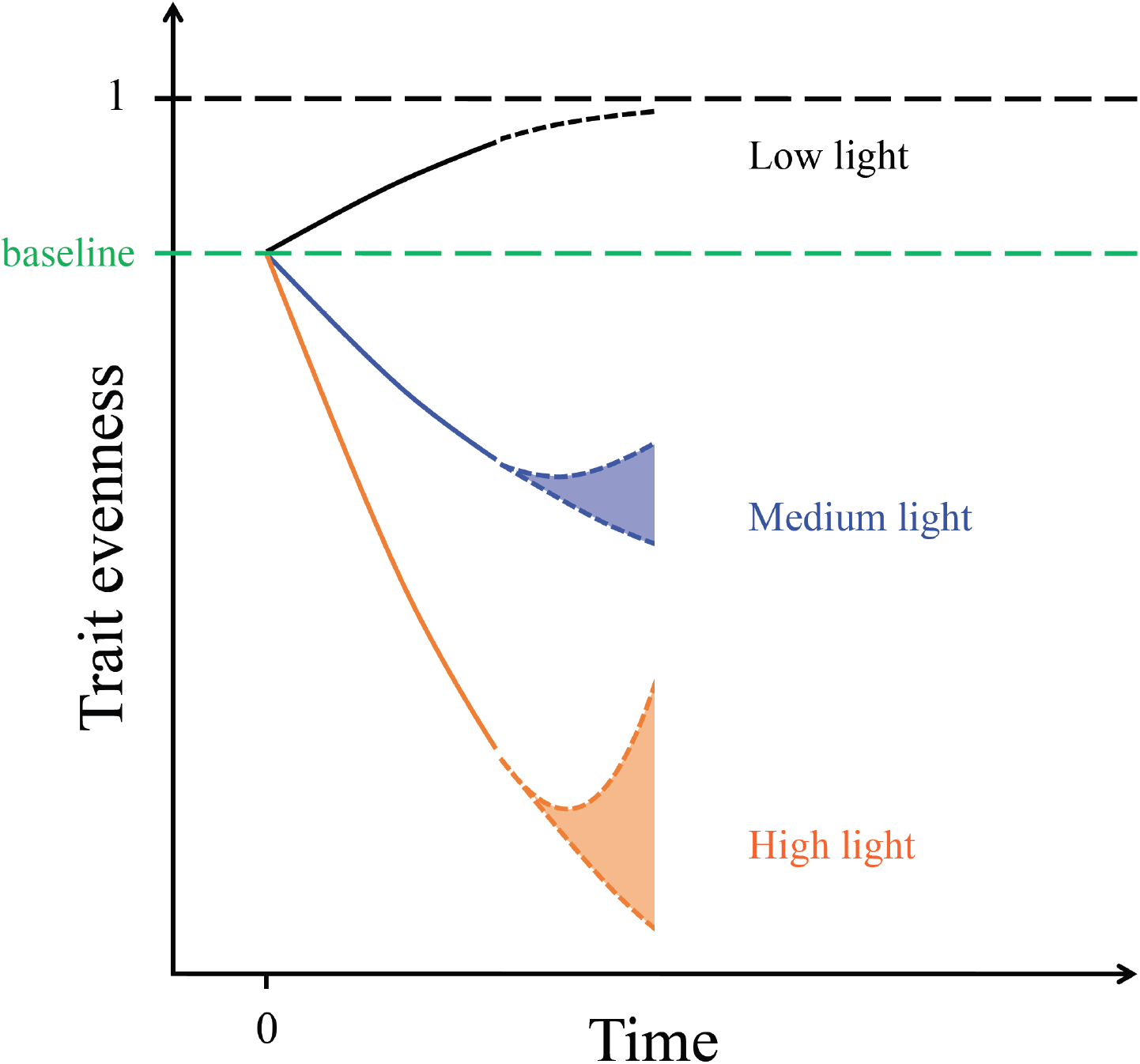
Expected change of trait evenness over time based on the hypothesis that light limitation and competition induce changes towards higher regularity in photosynthetic traits of individual phytoplankton cells. This likely results from the necessity to absorb wavelengths that are not used by competitors. The three depicted curves reflect different shifts in the light intensity regime compared to the time before the beginning of the experiment (decrease, moderate increase and strong increase). Dashed black=maximum possible value of trait evenness. Dashed green=baseline value of trait evenness, resulting from the maintenance of all cultures at the same light intensity prior to the experiment. We expect the response to be approximately linear at the beginning, whereas more uncertainty is associated with longer time scales, as indicated by coloured bands and shaded lines.

## Results

All results confirmed that decreasing light intensity induces an increase in the evenness of individual phytoplankton in multidimensional photosynthetic trait space.

Trait evenness (characterised by the TED index) decreased over time at all but the lowest light intensity, where it increased instead (Fig. 3). Trait evenness decreased more rapidly over time with increasing light intensity. Moreover, the rate of change in evenness over time was a nearly linear function of light intensity in both species (Fig. 4, top panels), consistent with our predictions. However, trait evenness decreased faster in *M. aeruginosa*, which also showed a much broader range of TED values compared to *P. subcapitata,* especially under direct light (Fig. 3; Fig. 4, bottom panels).

**Fig. 3.**
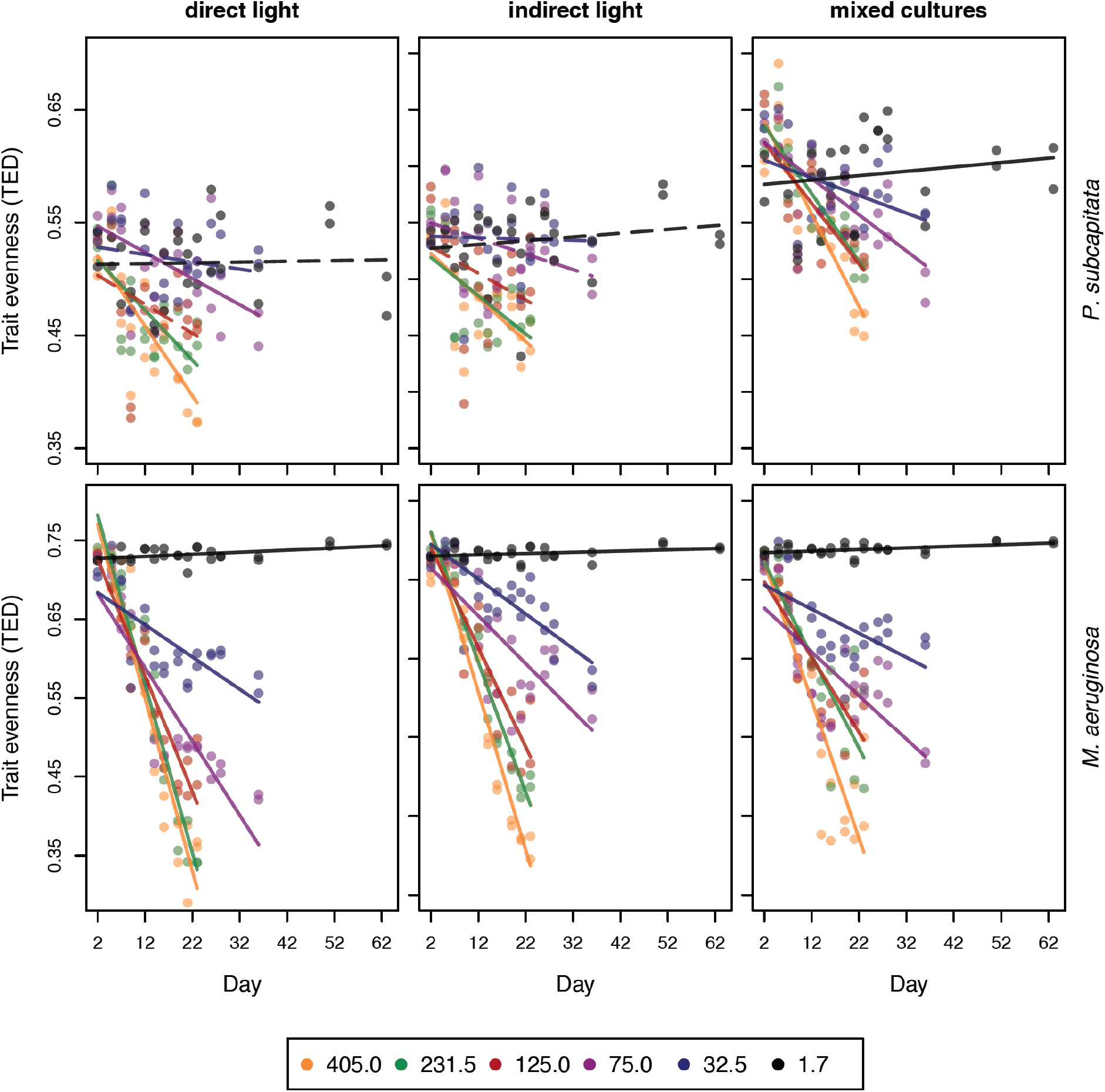
Change in trait evenness (TED index) over time. Lines represent independent linear model fits at each light level (in μE m^-2^ s^-1^) for each species (that we were able to distinguish also in mixed cultures, see Methods) and in each shading category. Dashed lines indicate non-significant relationships. Note that higher light intensity treatments stop earlier, as they reach carrying capacity faster and consequently show an accumulation of dead cells.

**Fig. 4.**
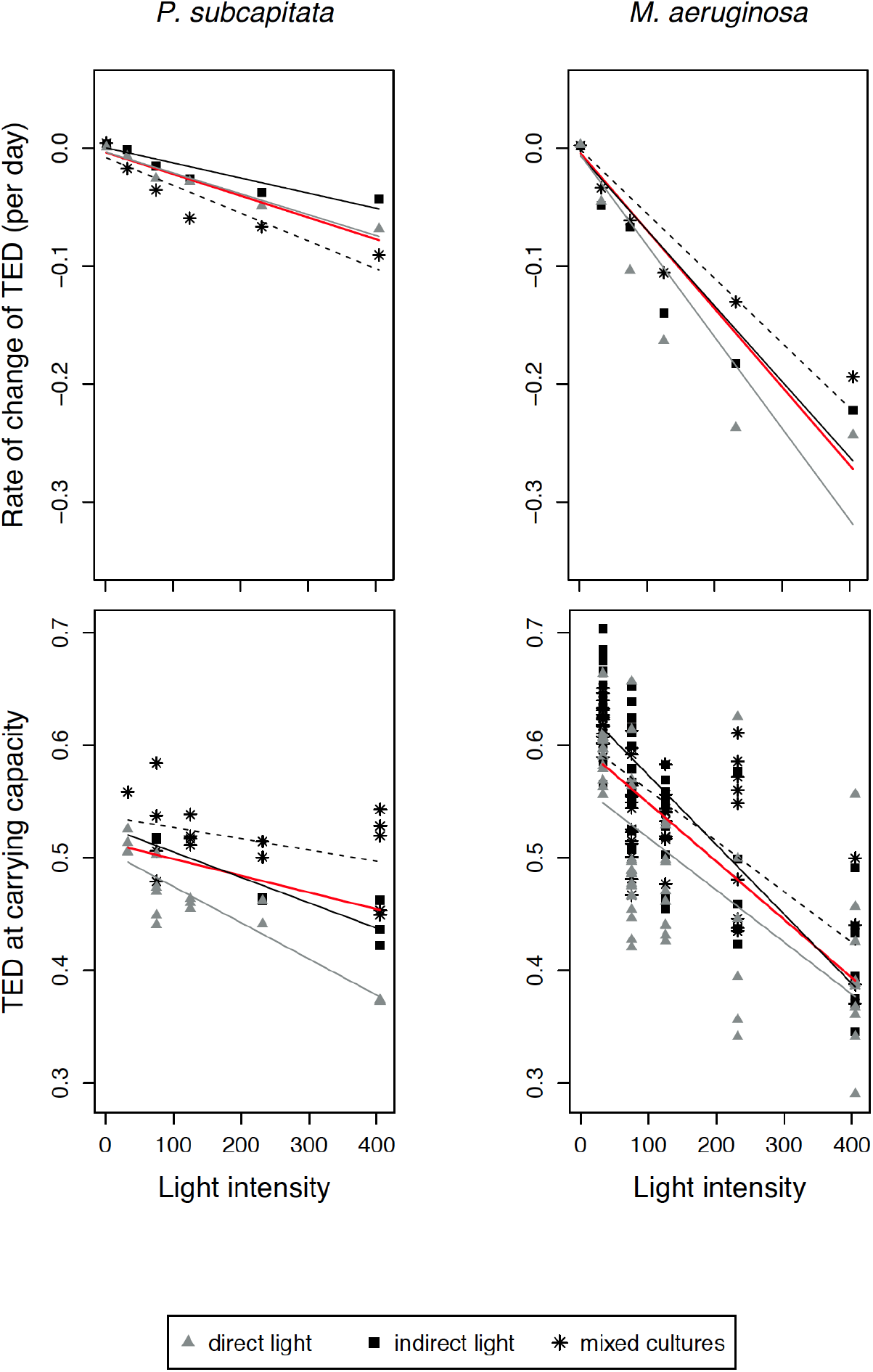
Relationship between light intensity and two response variables: slope estimate of each linear fit of TED change over time (top panels), and TED values at carrying capacity (bottom panels). Linear fit lines are indicated (dashed=mixed cultures, red=global fit). Note that in the bottom panel the number of points at each light intensity is not constant, because for each replicate we only included samples collected after reaching the carrying capacity (e.g. no samples in the lowest light treatment; Supplementary Fig. S2-S7).

Linear models of the rate of change in trait evenness explained between 75% and 93% of the variance (p<0.01) in both *P. subcapitata* and *M. aeruginosa* (Table 1A). Both *P. subcapitata* and *M. aeruginosa* displayed curves that appeared to be steeper under direct than under indirect light (Fig. 4, top panels). However, the slopes of the three shading levels (including mixed cultures) did not differ significantly, as indicated by the addition of the interaction between light intensity and shading in the global model *(P. subcapitata:* p = 0.10; *M. aeruginosa:* p = 0.41).

**Table 1.** Results of the linear models with light intensity (midpoint of the range in each light treatment) as predictor of two response variables: TED slope (=slope estimate of each linear fit of TED change over time; Fig. 3) (A), and TED values at carrying capacity (B). Regression estimates, standard errors, p-values and adjusted R^2^ are indicated. Global models include data points from monocultures (under direct and indirect light) and mixed cultures.

|  | *P. subcapitata* |  |  |  | *M. aeruginosa* |  |  |  |
| --- | --- | --- | --- | --- | --- | --- | --- | --- |
|  | Est.<br>[10<sup>-5</sup>] | SE<br>[10<sup>-5</sup>] | p-value | Adj. R<sup>2</sup> | Est.<br>[10<sup>-5</sup>] | SE<br>[10<sup>-5</sup>] | p-value | Adj. R<sup>2</sup> |
| **A) TED slope ~ light intensity** |  |  |  |  |  |  |  |  |
| global model | -18.4 | 2.5 | <0.001 | 0.75 | -66.4 | 6.1 | <0.001 | 0.87 |
| direct light | -17.7 | 2.2 | 0.001 | 0.93 | -77.7 | 14.7 | 0.006 | 0.84 |
| indirect light | -12.8 | 2.3 | 0.005 | 0.85 | -64.2 | 9.9 | 0.003 | 0.89 |
| mixed cultures | -23.6 | 4.6 | 0.007 | 0.84 | -54.9 | 7.1 | 0.001 | 0.92 |
| **B) TED carrying ~ light intensity** |  |  |  |  |  |  |  |  |
| global model | -14.9 | 4.7 | 0.003 | 0.19 | -51.7 | 3.8 | <0.001 | 0.52 |
| direct light | -32.2 | 4.4 | <0.001 | 0.78 | -46.3 | 6.5 | <0.001 | 0.44 |
| indirect light | -22.6 | 4.4 | 0.004 | 0.81 | -61.7 | 5.4 | <0.001 | 0.71 |
| mixed cultures | -10.0 | 5.8 | 0.11 | 0.12 | -45.0 | 6.1 | <0.001 | 0.49 |

Trait evenness at carrying capacity also decreased with increasing light intensity (Fig. 4, bottom panels); this decrease was statistically significant in all linear models with the exception of *P. subcapitata* in mixed cultures (Table 1B). Explained variance ranged between 12% and 81% (Table 1B). The slopes of the three shading levels were not significantly different for *M. aeruginosa* (interaction between light intensity and shading: p = 0.10). On the contrary, in *P. subcapitata*, trait evenness at carrying capacity decreased faster under direct than indirect light (Fig. 4, bottom panels), as shown by the significant interaction (p = 0.01) between light intensity and shading in the global model.

## Discussion

Our data show strong experimental evidence that 1) variation in resource availability leads to phenotypic trait combinations in phytoplankton that are detectable at both inter- and intraspecific levels and, 2) under conditions of limited resource (light) supply, the similarity between individual phytoplankton cells in terms of trait combinations tends to be minimised. Phytoplankton increased their phenotypic heterogeneity concomitantly reducing their trait similarity with hetero- or conspecific individuals. Specifically, our interpretation is that under conditions of light limitation, individual phytoplankton cells can reduce overlap in photosynthetic traits that support light acquisition, acquiring unexploited resources and thereby likely maximizing individual success. This causes individual phenotypes to partition evenly in multidimensional trait space (i.e. to show more regular distances among each other) when light intensity is low.

It has been suggested that considering several components of trait diversity help characterize complex changes in multidimensional trait space and effectively link phenotypic variation to community structural and functional properties (Mouillot *et al.* 2013; Fontana *et al.* 2016; Fontana *et al.* 2018). In this study, we used different light regimes as a way to manipulate resource limitation and competition, although one could argue that light limitation represents an environmental factor that typically constraints the number of viable trait combinations, thus reducing trait range and consequently increasing phenotypic similarity (Weiher & Keddy 1995; Kraft, Valencia & Ackerly 2008). Our observed patterns of an increase in even spacing of individuals in multidimensional trait space are not in contradiction with a possible, simultaneous contraction of the trait space covered (=reduction in trait richness). We note here that evenness in trait distribution, as measured by TED, is in fact a measure of trait regularity independent of the absolute distances between individual phenotypes (Fontana, Petchey & Pomati 2016) and reflects the tendency to maximise those distances given any trait range. Thus, even spacing of individuals is not necessarily related to the overall trait space covered (=trait richness). In our data, high TED was associated with high levels of trait richness (TOP index, Supplementary Fig. S1). The exception was represented by monocultures of *P. subcapitata,* which showed a non-significant relationship between TED and trait richness (Supplementary Fig. S1), probably influenced by a lower diversity in pigment FL-related traits in this species compared to *M. aeruginosa* (more constrained TOP and TED values in Supplementary Fig. S1).

Overall, during the experiment, trait evenness decreased faster and showed a broader range of values in *M. aeruginosa* (Fig. 3; Fig. 4) compared to *P. subcapitata. M. aeruginosa* also outperformed *P. subcapitata* in terms of growth in the mixed cultures, under all light levels (Fig. S2-S7). These results suggest that the cyanobacterium is capable of a higher photosynthetic trait plasticity compared to the green alga (Tandeau de Marsac 1991; Stomp *et al.* 2004; Mantzouki *et al.* 2016), and benefits from a faster growth possibly due to a faster metabolism and a broader array of available pigments (Richardson, Beardall & Raven 1983; Glazer 1984). The importance of phenotypic plasticity (adjustment of pigment ratios to the prevailing light spectrum) in determining the outcome of competition for light has been demonstrated before in phytoplankton, but only at the interspecific level (Stomp *et al.* 2008). Most likely in our case, phenotypic changes can be ascribed to plasticity at the very short time scale (days) after treatments were applied (Fig. 3), while selection and evolution could have played a role in the observed patterns at the end of the experiment (Ellner, Geber & Hairston 2011). More work will be needed in the future to understand the relative importance of invoked mechanisms such as plasticity, evolution and competitive differences in shaping the patterns observed here. Similarly, the generality of our findings across other taxonomic groups, morphological and physiological traits or resource gradients requires assessment.

This study however supports the hypothesis that the regularity in the distribution of individuals along multiple resource acquisition axes increases with decreasing resource availability. Our results reinforce previous findings that resource limitation increases phenotypic heterogeneity in microbes (Schreiber *et al.* 2016), and indicate that a highly restrictive abiotic environment (i.e. low light) does not necessarily select for a reduced number of trait combinations, as often assumed in ecology (the environmental filtering hypothesis) (Kraft *et al.* 2015). Instead of convergence towards an optimal phenotype, survival under low resource conditions in our experiment required cells to find new (unique) strategies, thus maximising at the same time phenotypic distances in trait space. This finding is consistent with the hypothesis that high evenness in distribution of individuals in trait space emerges in conditions in which it is advantageous to escape or minimise competition. On the contrary, when resources are not limiting, organisms may converge towards few phenotypic profiles that allow the fastest growth rate, as indicated in our experiment by declining regularity in phytoplankton trait distributions and amount of trait space covered. In summary, we report novel empirical insights into how resource availability can shape phenotypes in a competitive environment, and demonstrate that conspecific and heterospecific phytoplankton cells tend to differentiate their pigment profile under conditions of low light intensity. Our results represent a step forward in elucidating the mechanisms that maintain individual-level coexistence in populations and communities over resource gradients.

## Methods

### Experimental conditions and culture acclimation

In a climate-controlled room with constant temperature of 20°C, we set up six different light regimes. Osram T8 L De Luxe 36W 954 G13 Lumilux lights were placed under Plexiglas supports of varying height. The support elevation and several meshes (black and white) were used to vary light intensity (more details on experimental setup in Supplementary Table S1). The ranges of the resulting light intensities (in μE.m^-2^.s^-1^) at the culture bottles over multiple measurements are presented here: 380-430 (midpoint=405), 173-290 (231.5), 95-155 (125), 60-90 (75), 25-40 (32.5), 0.4-3 (1.7). All treatments followed a 14:10 hours light-dark cycle.

*P. subcapitata* strain SAG 12.81 (from the SAG Culture Collection of Algae, University of Göttingen, Germany) and *M. aeruginosa* strain PCC 7806 (toxic wild type obtained from Brett A. Neilan, UNSW, Sydney, Australia) were maintained as batch cultures in a separate culturing room under identical environmental conditions. Cultures were maintained in exponential growth phase at 20°C and a light intensity of approximately 20-25 μE m^-2^ s^-1^ (intermediate between the two lowest light intensity treatments). The initial trait evenness of our test cultures was therefore supposed to be similar at the beginning of the experiment (Fig. 2, dashed green baseline). Replicated cultures were transferred to the different experimental light regimes 10 days before starting the experiment.

We diluted with WC-medium the content of the Erlenmeyer flasks (batch cultures) into cell culture bottles (Faust Lab Science, product number TPP90301) closed with filter caps that allow gas transfer. We applied a 1:20 dilution of the original cultures for all but the two lowest light intensities (1:10 in this case, as we expected a much slower growth under low light regimes), with mixed cultures having equal volumes of the two species, and obtained a final volume of 200 mL in each cell culture bottle.

### Experimental design

We obtained a total of six units in every light treatment compartment: cultures placed on top of each other *(P. subcapitata* above – and shaded by – *M. aeruginosa,* plus *M. aeruginosa* above *P. subcapitata*, two replicates each) and mixed cultures (two replicates) (Fig. 1). In this way, monocultures of *P. subcapitata* and *M. aeruginosa* were subjected to both direct and indirect light (cell culture bottles were narrow and so did not totally block incoming light), while mixed cultures always experienced direct light. The position of these units within the light treatment compartments was randomised and shuffled regularly. The experimental design is illustrated in Fig. 1.

### Culture maintenance and sampling protocol

Each culture was manually shaked once a day from Monday to Friday to avoid the formation of a biofilm on the flasks’ bottom. The three highest light intensity treatments (405, 231.5 and 125 μE m^-2^ s^-1^) were sampled 10 times between the 9^th^ of March (start of the experiment) and the 1^st^ of April 2016 (every 2-3 days for 24 days), when they had already reached carrying capacity and showed an accumulation of dead cells (based on colour change). The three lowest light treatments (75, 32.5 and 1.7 μE m^-2^ s^-1^) were additionally sampled on the 4^th^, 6^th^ and 14^th^ of April, and the very lowest also on the 29^th^ of April and on the 11^th^ of May. These additional sampling dates allowed all treatments but the lowest to reach carrying capacity (as defined in the paragraph “Logistic growth and carrying capacity”). At each sampling date we collected 1.5 mL from all cultures in Eppendorf tubes and fixed it with 0.01% paraformaldehyde and 0.1% glutaraldehyde for SFC analyses.

### SFC measurements

We characterized the fluorescence profile (related to light acquisition strategy) of individual cells in each population / community using SFC. The instrument we used (www.cytobuoy.com, Woerden, The Netherlands) is able to capture scattering and pigment fluorescence of algal cells in a time resolved mode (scanning) (Dubelaar, Geerders & Jonker 2004; Pomati *et al.* 2013). The instrument is specifically designed for high resolution scanning of freshwater phytoplankton pigment fluorescence emission. The scanning flow-cytometer was equipped with two laser beams (coherent solid-state sapphire) with excitation wavelengths of 488 nm (blue light) and 635 nm (red light), which allow to excite all algal main and accessory pigments. Four detectors captured emitted pigment fluorescence in the red (677–700 nm, mainly chlorophyll-a), orange (650–677 nm, for phycocyanin), yellow (590–620 nm, for phycoerythrin), and green (550–570 nm, for carotenoids) ranges. Data acquisition during this study was triggered by sideward scattering (SWS signal, trigger threshold 30 mV), and the flow speed was 1.05 μL s^-1^. We measured approximately between 6’000 and 99’000 cells depending on cell densities.

Applying a clustering algorithm *(flowPeaks* package, R-Core-Team 2013) to a subset of 100’000 particles built by randomly taking the same amount of particles from each experimental sample, we were able to define distinct clusters of organic debris / suspended solids (characterized by low FL), *P. subcapitata* and *M. aeruginosa* (the second having a higher orange to red fluorescence ratio). Using this sample dataset, we trained a random forest classifying algorithm (Breiman 2001; Liaw & Wiener 2002) and assigned every particle in the whole dataset to one of the three abovementioned categories (Thomas *et al.* 2018).

### Individual-based trait distribution

The regularity in the distribution of individual cells in multidimensional trait space was quantified for all samples using the TED index (Fontana, Petchey & Pomati 2016), which compares pairwise distances between individuals with those in a perfectly even reference distribution. For the calculation of this multivariate index four SFC-derived traits were used, which are highly relevant in competition for light: total fluorescence red, orange, yellow and green. These traits are associated with pigment quantity and activity, and are therefore indicators of relative investment of individual cells in capturing different portions of the light spectrum (Stomp *et al.* 2007; Pomati *et al.* 2013; Fontana *et al.* 2018). A subset of 5’000 randomly selected cells was used to calculate TED. For mixed cultures, we additionally calculated species-specific TED indices with a random subset of 1’000 cells for each species. To test the relationship between trait evenness (TED) and trait richness, we also calculated the TOP index (Fontana, Petchey & Pomati 2016) of the same subsets of cells used for TED. TOP is a measure of trait richness that takes into account all individual phenotypes (including intermediate ones), and thus does not simply represent a multidimensional range but rather an estimation of the trait space effectively covered by individuals (Fontana, Petchey & Pomati 2016).

### Logistic growth and carrying capacity

Cell counts from SFC were used to derive cell density of each sample. We fitted a logistic growth curve to cell density data from each of the replicates using the function *gcFitModel (grofit* package, R-Core-Team 2013) to obtain carrying capacity estimates (details in Supplementary Table S2). R^2^ values, calculated regressing observed against fitted values, ranged from 0.845 and 0.996 (Supplementary Table S2). In each replicated growth curve (Supplementary Fig. S2-S7) we identified the first time point with density equal to or greater than the carrying capacity estimate. We defined all time points after reaching carrying capacity as the period of maximum competition for resources among individuals, which allowed a meaningful comparison of trait evenness across treatments, independent of the different temporal trajectories, by controlling for the effect of population-specific growth over time.

### Statistical analyses

Our experiment covered a time interval insufficient to detect clear curvilinear trends, and we expected the response to be approximately linear at the beginning (Fig. 2). Therefore, we used simple linear regressions as the most parsimonious common approach for all analyses (after checking model assumptions visually).

To test whether even spacing of individuals across photosynthetic trait axes represented a response to light limitation, we performed two different analyses considering *P. subcapitata* and *M. aeruginosa* separately. First, we fitted linear regressions for each combination “light treatment × shading” separately (Fig. 3), with date of sampling as predictor and TED as response variable. All slope estimates were then included as response variable in linear models, using least squares weighted by the inverse of slope standard errors (to account for uncertainty in slope estimates) and light intensity midpoint as predictor (mean value of minimum and maximum measured for each treatment). Second, using the time points at carrying capacity detected in replicates (as defined in the paragraph “Logistic growth and carrying capacity”), we fitted linear models to test the relationship between light intensity midpoint (predictor) and TED at carrying capacity. Replicates of the lowest light treatment did not reach carrying capacity and consequently were omitted from these linear models. This analysis aimed at testing the influence of light intensity on TED by correcting for its temporal change, which varied among different cultures. Analysis of covariance (ANCOVA) was used to test whether shading levels (“direct light”, “indirect light” and “mixed cultures”) influenced the effect of light intensity on the rate of change of TED and TED at carrying capacity in the two above-mentioned analyses. To this end, using the function *aov* (R-Core-Team 2013) we included the interaction between shading and light intensity in the global models. We calculated type-III analysis-of-variance tables (orthogonal contrasts setting) with the function *Anova* (*car* package, R-Core-Team 2013) to obtain F-tests of the explanatory variables.

## Acknowledgements

This research is supported by the Swiss National Science Foundation Projects 31003A_144053 and CRSII2_147654. We thank R. Stegmayer for his help in lab work, H. Penson for maintaining algal cultures, and M. Kehoe for the inspiring discussions. The Centre for Ocean Life is a Villum Kahn Rasmussen Centre of Excellence funded by the Villum Foundation. The authors dedicate this work to the memory of M. Stomp.

